# Comparison of Different Approaches to Single Cell RNA Sequencing of Cancer Associated Fibroblasts

**DOI:** 10.1101/2024.04.29.591011

**Authors:** Heng-Chung Kung, Michael Loycano, Lei Zheng, Sophia Y. Chen, Jacquelyn W. Zimmerman

## Abstract

**Background:** Pancreatic ductal adenocarcinoma (PDAC) is a highly aggressive disease with a poor prognosis. PDAC has a high propensity for metastasis, particularly to the lungs and liver. Cancer associated fibroblasts (CAFs) represent a major stromal component of PDAC with both tumor-promoting and restraining properties. Of note, CAFs play a significant role in the creation of an immunosuppressive tumor microenvironment (TME) and the metastasis of PDAC. Studies have demonstrated functional heterogeneity among different subpopulations of CAFs, highlighting the need to identify specific subpopulations when targeting CAFs.

**Methods:** The orthotopic model was used for both KPC-4545 and KPC-3403 cell lines, which were derived from the primary tumors of KPC mice with liver metastases and lung metastases only, respectively. In brief, 2x10^6^ KPC cells were injected subcutaneously into the flanks of synergic female C57BI6 mice. Tumors were harvested and cut into 2-3 mm^3^ pieces before being implanted into the pancreas of new 6–8-week-old syngeneic female C57Bl/6 mice. Murine orthotopic tumors were dissected, mechanically and enzymatically processed with Miltenyi Tumor Dissociation Kit (Miltenyi Biotec) thirteen days after tumor implantation. Samples were filtered with a 100 µm strainer, washed with T cell media, and centrifuged twice.

Two different samples underwent single cell RNA-sequencing (scRNA-seq) for each cell line: an unenriched sample, which represents all cells following dissociation of the tumor, and a CAF-enriched sample. To further obtain the CAF-enriched sample, cells were then stained with CD45-AF657 (BioLegend clone 30-F11, 1:20), CD31-AF647 (BioLegend clone 390, 1:20), EPCAM-AF647 (BioLegend, clone G8.8, 1:20), and TER119-AF647 (BioLegend clone TER-119 1:20) for 30 minutes on ice. After two washes, CD45-, CD31-, EPCAM-, and TER119-negative cells, representing the CAF-enriched fraction, were obtained via cell sorting. scRNA-seq of both the unenriched and CAF-enriched fractions were performed using 10X Chromium microfluidic chips and data was analyzed using CellRanger v6.1.1, mm10 transcriptome reference, and 10X Loupe Browser.

**Results:** We found that scRNA-seq of the unenriched whole tumor showed only one cluster of CAFs for both cells lines, making it difficult for studying CAF heterogeneity. Enriching for CAFs prior to scRNA-seq allowed for better capture of CAFs and provided more granularity on CAF heterogeneity for both KPC-4545 and KPC-3403.

**Conclusions:** While enrichment provides more information on CAF heterogeneity, the process results in the loss of other cells within the TME. The need to capture CAF heterogeneity while studying cell-cell interaction between CAFs and other cells within the TME and identifying how distinct CAF populations respond differently to treatment warrants the use of other methods such as single-nuclear RNA-seq.

## INTRODUCTION

Pancreatic ductal adenocarcinoma (PDAC) is one of the most aggressive and lethal malignancies due to its high propensity for metastasis and high resistance to existing therapies (1, 2). With a 5-year survival rate of 13%, PDAC is projected to be the second leading cause of cancer-related deaths in 2030 (3). Nearly 50% of patients present with metastatic disease at the time of diagnosis, with the liver, lungs, and peritoneum being the most common sites of metastasis (4, 5). Our previous clinical observations have demonstrated differential outcomes of PDAC based on different metastatic sites, with lung metastasis performing better compared to liver metastasis (6). However, the mechanisms of site-specific metastasis in PDAC are not fully understood.

Contributing to the poor outcomes of PDAC is its immunosuppressive and dense tumor microenvironment (TME), featuring a wide range of infiltrating cell types such as mesenchymal cells and different immune cell subtypes (7). Notably, cancer associated fibroblasts (CAFs) represent a major stromal component of the PDAC TME, with studies demonstrating both its tumor-promoting and tumor restraining properties (8). Recent research has explored the heterogeneity of CAF phenotypes within the PDAC TME, identifying functionally distinct populations of CAFs (9). Myofibroblastic CAFs (myCAFs) and inflammatory CAFs (iCAFs) have been found in both human and mice tumors while antigen-presenting CAFs (apCAFs) have only been identified in mice tumors (10). Different CAF populations have been shown to engage in complex cell-cell interaction and signaling with malignant cells and other stromal cells (11). Particularly, CAFs are known to promote metastasis by mediating epithelial-mesenchymal transition (EMT) (12).

In order to better understand the mechanisms behind the site-specific metastasis in PDAC, our group previously established liver-specific and lung-specific KPC PDAC tumor cell lines (13). Using RNA whole-transcriptome sequencing, we found that mouse mesenchymal stem cells (moMSCs) co-cultured with liver-metastasis KPC cells presented with a more homogenous CAF phenotype while moMSCs co-cultured with the lung-metastasis KPC cells retained their heterogenous CAF phenotypes (13). Over the past few years, single cell RNA-sequencing (scRNA-seq) has allowed for more in-depth study on the PDAC TME, enabling researchers to not only identify discrete cell-type populations and phenotypes, but to also decode complex cell-cell interaction between different components of the TME. Here we attempt to study the stromal composition and CAF phenotypes of our site-specific metastasis PDAC murine model using scRNA-seq (14).

## METHODS

### Cell lines

The KPC (LSL-Kras (G12D/+); LSL-Trp53 (R172H/+); Pdx-1-Cre) tumor cell line is a previously established PDAC cell line from a KPC transgenic mouse model in a C57BL/6 background (15). The KPC-4545 cell line was derived from the primary tumor of a KPC mouse with liver metastasis only while the KPC-3403 cell line was derived from the primary tumor of a KPC mouse with lung metastasis only as previously described (13). KPC cell lines were cultured in RPMI 1640 media (Life Technologies) with 10% fetal bovine serum (FBS, Benchmark), 1% MEM-NEAA (minimum essential medium-non-essential amino acids, Life Technologies), 1% penicillin/streptomycin (Life Technologies), 1% sodium pyruvate (Sigma), and 1% L-glutamine (Life Technologies). Tumor samples were harvested and processed in T-cell media, consisting of RPMI 1640 (Life Technologies) supplemented with 10% heat-inactivated FBS (Benchmark), 1% penicillin/streptomycin (Life Technologies), 1% HEPES (Life Technologies), 1% MEM Non-Essential Amino Acids Solution (Life Technologies), 1% L-glutamine (Life Technologies), and 0.05% 2-mercaptoethanol (Sigma-Aldrich).

### Mice Studies

C57BI6 mice (6-8 weeks) were purchased from Jackson Laboratories, and all mice experimental protocols were performed in accordance with the Johns Hopkins University Institutional Animal Care and Use Committee (IACUC) guidelines. The KPC orthotopic models were established as previously described for both the KPC-4545 and KPC-3403 tumor cell lines. In brief, 2x10^6^ KPC cells were injected subcutaneously into the flanks of synergic female C57BI6 mice. After 1-2 weeks, the subcutaneous tumors were harvested and cut into 2-3 mm^3^ pieces. New 6–8-week-old syngeneic female C57Bl/6 mice were anesthetized. A left subcostal incision was made in the abdomen to access the body and tail of the pancreas. Using microscissors, a small pocket was prepared in the middle of the pancreas. One 2-3 mm^3^ piece of the subcutaneous tumor was implanted into the small pocket and secured with 7-0 Prolene suture. The peritoneal and skin were closed in two layers using 4-0 sutures.

### Cell Processing

Murine orthotopic pancreatic tumors were dissected and collected on day 13 after tumor implantation for scRNA-seq processing. Tumors were mechanically and enzymatically processed with the mouse Miltenyi Tumor Dissociation Kit (Miltenyi Biotec) and gentleMACS Octo Dissociator (Miltenyi Biotec). Samples were then filtered through 100 µm strainer, which was then washed with 10 mL T-cell media. Cell suspensions were centrifuged at 1500 RPM for 5 min. Cell pellets were resuspended in 10 mL T-cell media, filtered through 100 µm strainer, and centrifuged at 1500 RPM for 5 minutes again.

For fibroblast enrichment, cells were resuspended with PBS and blocked with anti-mouse Fc antibody (BD Biosciences) in FACS buffer for 10 minutes on ice. Cells were then stained with CD45-AF657 (BioLegend clone 30-F11, 1:20), CD31-AF647 (BioLegend clone 390, 1:20), EPCAM-AF647 (BioLegend, clone G8.8, 1:20), and TER119-AF647 (BioLegend clone TER-119 1:20) for 30 minutes on ice. Single cell suspension was mixed every 10 minutes. Cells were washed with PBS twice, resuspended at 5-10 million cells/mL in T-cell media, and filtered through 35 µm flow cytometry tubes (Falcon 352235). Sorting for CD45-, CD31-, EPCAM-, and TER119-negative cells was performed to obtain the fibroblast enriched fraction. Both the unenriched and enriched samples were washed and resuspended to cell concentration of 700-1200 cells/µL in PBS + 0.1% BSA for loading onto 10X Chromium microfluidic chips.

### Single cell RNA-sequencing processing and analysis

Single-cell capture, barcoding, and library preparation was performed using 10X Chromium Single Cell 5’ R2 chemistry according to the manufacturer’s protocol (#CG000207). CellRanger version 6.1.1 (10X Genomics) was used to convert Illumina base calls into FASTQ files and align FASTQ to the mm10 transcriptome reference. All scRNA-seq data was analyzed via the Loup Browser (10X Genomics). Cells were clustered with the graph-based clustering method provided by the Loupe Browser and was represented through uniform manifold approximation and projection (UMAP). Clusters were identified with cell-type specific marker genes curated from literature, and heatmap of labeled clusters and top expressed marker genes were created with the heatmap function on the Loupe Browser (16, 17). Violin plots for CAF phenotype were generated with the Loupe Browser using published marker genes for each CAF phenotype (**Table S1**).

## RESULTS

Within the KPC-4545 unenriched sample, a total of 836 cells were clustered into five clusters, representing a mixture of acinar and erythroid cells; a mixture of ductal cells, CAFs, and granulocytes; B/Plasma; T cell/natural killer (T/NK) cells; erythroid cells (**Figure 1A, 1B; Table S2**). Similarly, in the KPC-3403 unenriched sample, a total of 1,831 cells were clustered into eight clusters, representing CAFs; a mixture of erythroid and acinar cells; erythroid cells (Erythroid 1 and Erythroid 2); T/NK cells; granulocytes; ductal cells; B/Plasma cells (**Figure 1C, 1D; Table S3**). CAFs accounted for less than 21.4% and approximately 16.2% of all cells in the KPC-4545 and KPC-3403 unenriched samples respectively.

**Figure 1.**
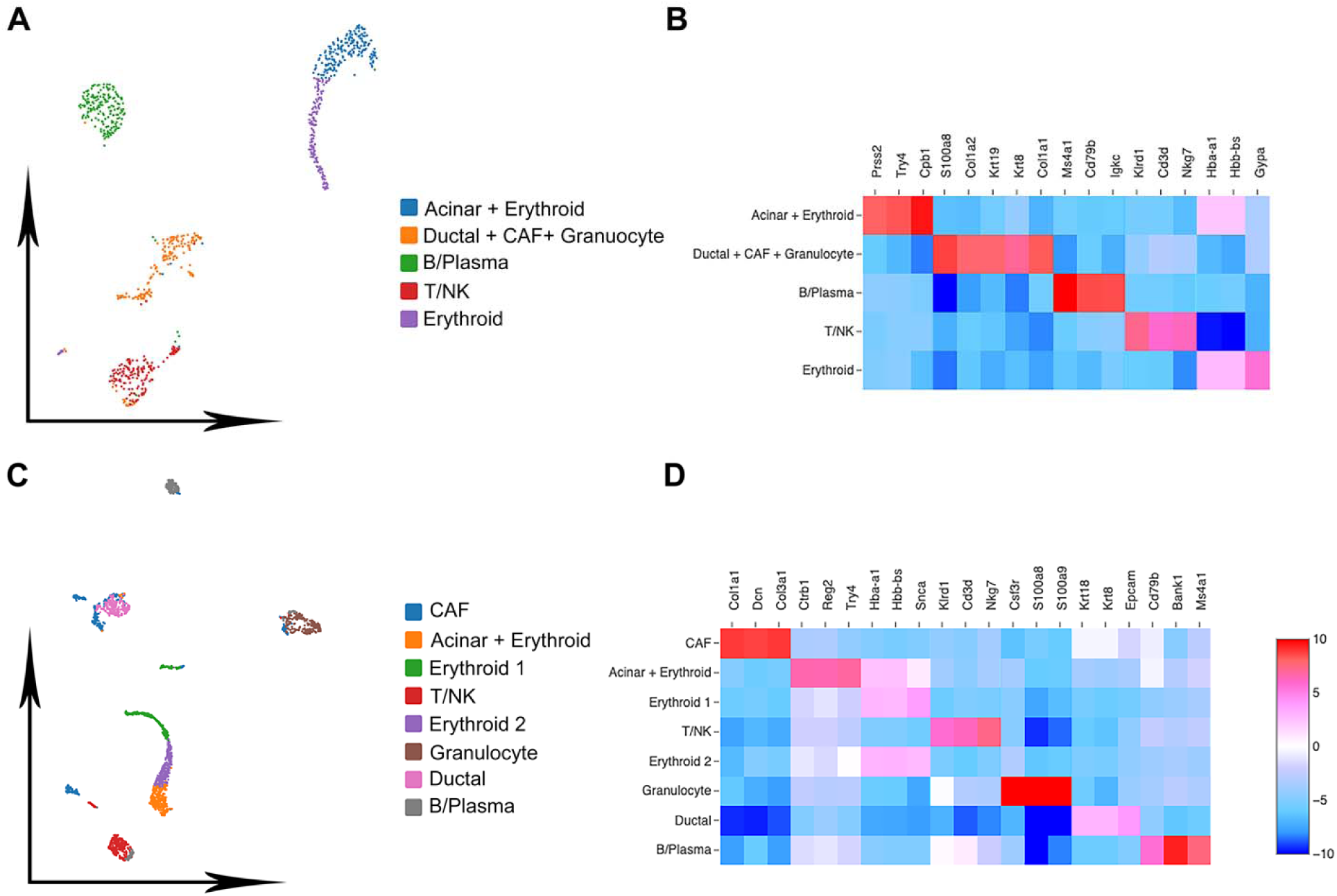
Single cell RNA sequencing of unenriched single cell dissociates from whole tumors for KPC-4545 and KPC-3403. **(A-B)** KPC-4545 unenriched sample UMAP **(A)** and cell type marker genes **(B)**. **(C-D)** KPC-3403 unenriched sample UMAP **(A)** and cell type marker genes **(B)**.

Following CAF enrichment via negative-FACS selection and scRNA-seq processing, three clusters were removed due to enrichment in mitochondrial genes representative of dying cells, resulting in a total of 2,028 cells in five clusters in the KPC-4545 enriched samples (**Figure 2A**). Based on marker genes curated from literature (**Supplementary Table 1**), all five clusters displayed strong panCAF marker expression (**Figure 2B**), confirming their identities as CAFs.

**Figure 2.**
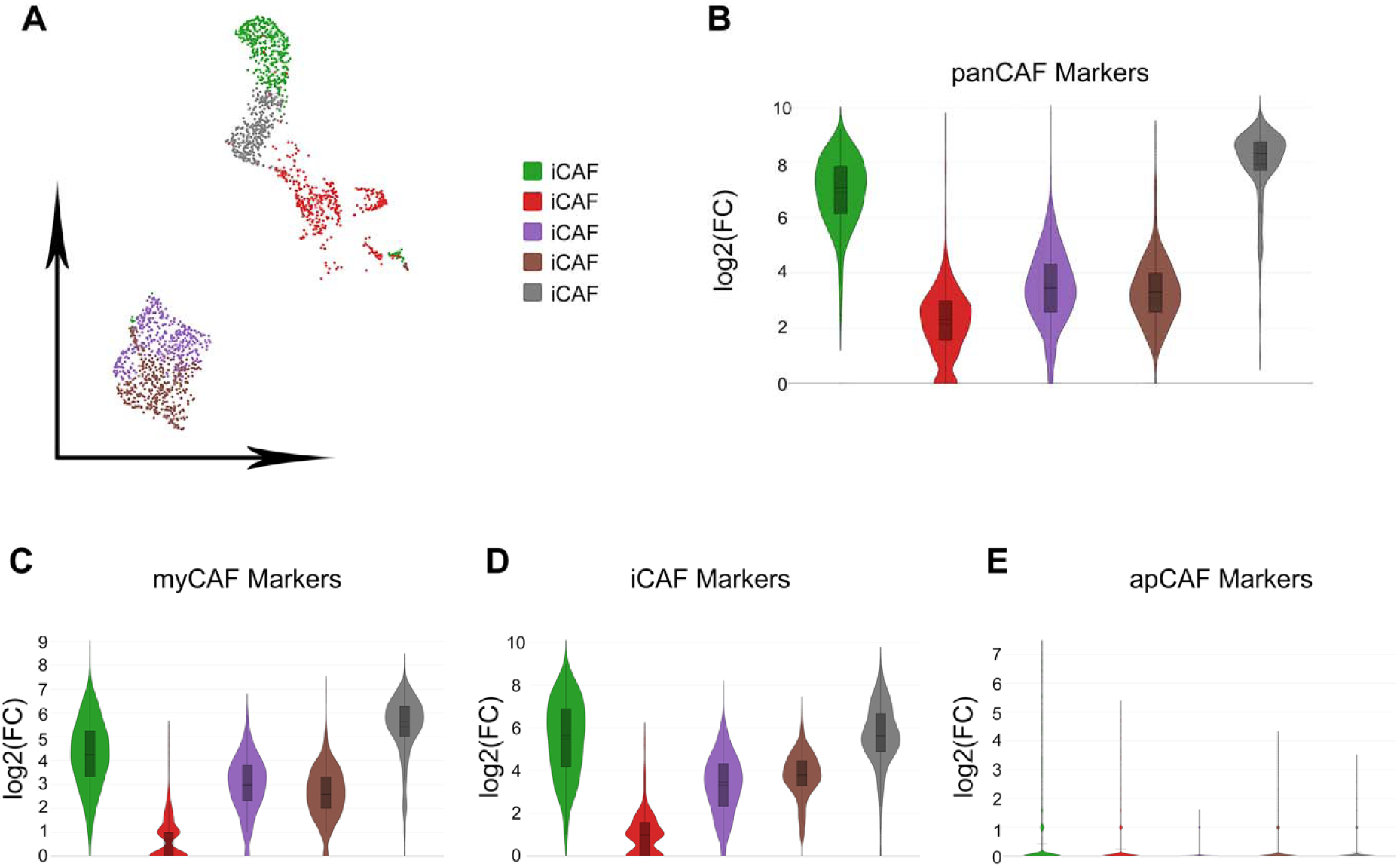
Single cell RNA sequencing of KPC-4545 CAF-enriched sample. (A) UMAP of KPC-4545 CAF-enriched sample. (B-E) Marker gene expressions for panCAF (B), myCAF (C), iCAF (D), and apCAF (E) in KPC-4545 CAF-enriched sample.

However, while labeled as iCAFs, all five clusters displayed relatively equal expression of myCAF and iCAF markers (**Figure 2C, 2D**), suggesting the presence of CAF populations with a more homogeneous CAF phenotype within the KPC-4545 TME. Expression of apCAF markers in the KPC-4545 enriched sample was negligible (**Figure 2E**).

In the KPC-3403 enriched sample, one cluster was removed due to enrichment in mitochondrial genes, resulting in 2,753 cells in eight clusters (**Figure 3A**). All clusters except for one expressed panCAF markers (**Figure 3B**), which we identified as myeloid cells. Contrary to the homogenous CAF phenotype observed in the KPC-4545 enriched sample, CAFs in the KPC-3403 adopted more heterogenous phenotypes, with the presence of myCAFs clusters, two iCAFs clusters, three apCAFs clusters as identified by marker gene expression (**Figure 3C-E**).

**Figure 3.**
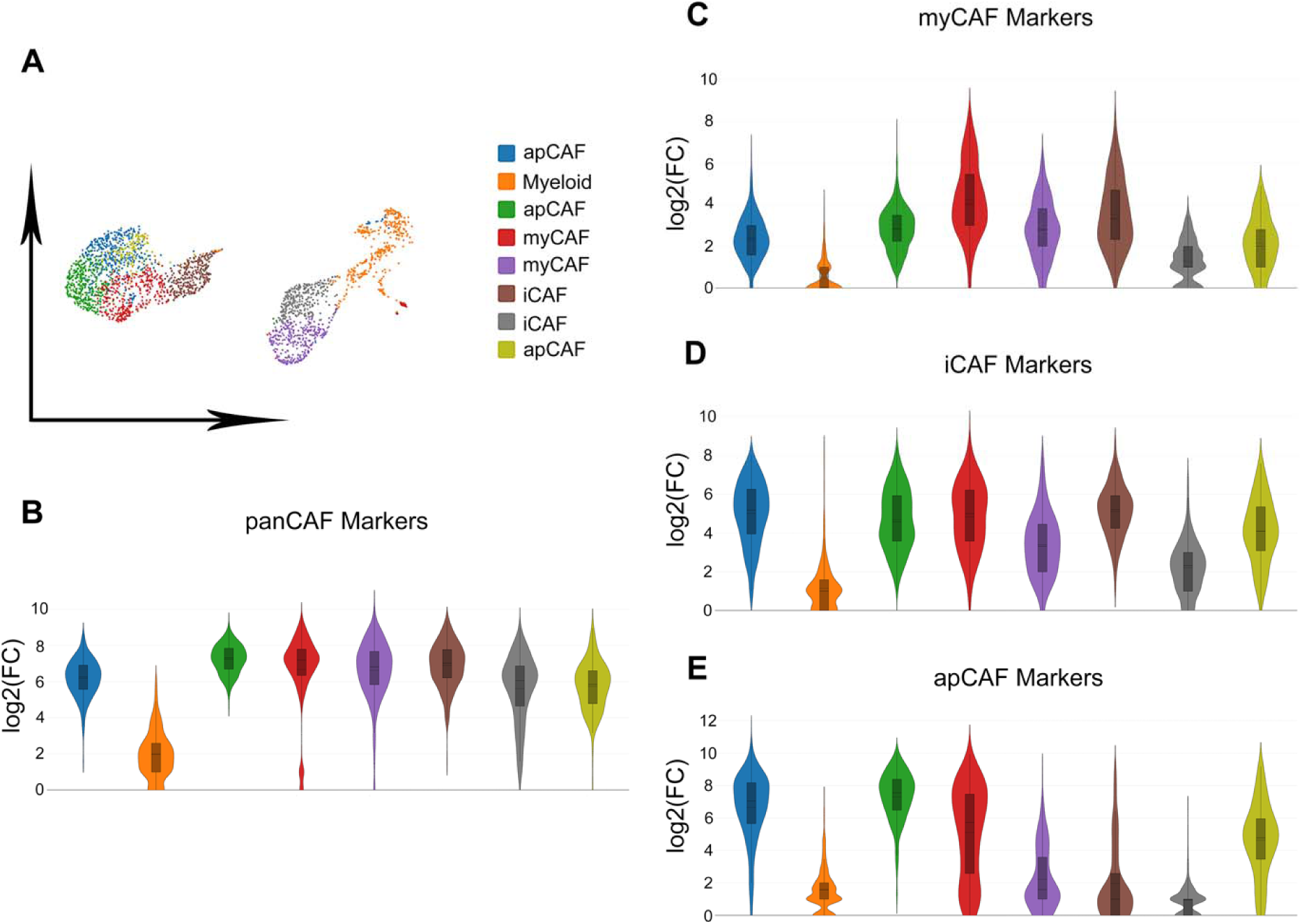
Single cell RNA sequencing of KPC-3403 CAF-enriched sample. **(A)** UMAP of KPC-3403 CAF-enriched sample. **(B-E)** Marker gene expressions for panCAF **(B)**, myCAF **(C)**, iCAF **(D)**, and apCAF **(E)** in KPC-3403 CAF-enriched sample.

## DISCUSSION

In this study, we tested the feasibility of investigating CAF phenotypes using scRNA-seq of orthotopic KPC PDAC model by using unenriched (whole tumor) and CAF-enriched samples. We found that CAFs only represented a fraction of cells in unenriched samples without sufficient granularity for identifying different subpopulations/phenotypes. Enriching for CAFs allowed for higher resolution of CAF phenotypes, but at the sacrifice of profiling other cells.

Previous studies on PDAC in the past have primarily focused on the development of anti-cancer therapies targeting the epithelial compartment (18). However, research over the past decade has increasingly shown that stromal components of PDAC play a major role in the tumorigenesis and progression, particularly CAFs (19). On one hand, numerous studies have demonstrated the protumor properties of CAFs. As the main source of extracellular matrix (ECM), CAFs secrete a wide range of ECM proteins, including hyaluronan and collagen (20).

This can lead to the “hardening” of the ECM, which hinders the delivery of drugs and infiltration of immune cells to the cancer cells encased by the ECM (21, 22). Furthermore, CAFs inhibit cytotoxic T cell activity in the TME by secreting TGFβ, IL-6, and CXCL12 (23, 24); recruit immunosuppressive populations such as neutrophils (25); and promote the polarization of M2 macrophages to establish an immunosuppressive TME (26).

Given these protumor mechanisms, much work has been put into targeting CAFs as possible treatment for PDAC, albeit with mixed results. Particularly, studies depleting alpha-smooth muscle actin (αSMA) positive CAFs or inhibiting the Hedgehog (Hh) pathway have reported reduced survival and worse outcomes in both preclinical models and clinical studies (27–30). These results suggest the potential tumor-restraining role of CAFs and the presence of different subtypes of CAFs with heterogenous functions (31). This heterogeneity warrants a more in depth understanding of the functions of different CAF phenotypes for a more targeted approach against CAFs in the treatment of PDAC.

However, the study of CAFs has proven difficult due to the lack of precise fibroblast-specific markers (32). The absence of exclusive cell marker is partially due to CAFs’ wide range of possible cellular precursors, including quiescent pancreatic stellate cells (PSCs), residential fibroblasts, tumor-infiltrating mesenchymal stem cells (MSCs), and mesothelial cells (33).

Different CAF phenotypes were first observed through immunofluorescence studies, and more subtypes and associated markers have been identified with the use of multi-color flow cytometry and immunohistochemical studies (18, 34, 35). Over the past ten years, scRNA-seq technology has allowed for the investigation of CAF phenotypes through their transcriptomic profiles at a single cell level, allowing researchers to further unravel CAF heterogeneity in the TME by discovering new markers and understanding subpopulation functional states based on their gene expression (10, 36–39). This has resulted in a spectrum of markers for CAFs, including intracellular markers (VIM, FSP-1, αSMA), extracellular markers (LUM, DCN, COL1A1, COL1A2), and surface markers (FAP, PDPN, PDGFRα, PDGFRβ) (18). Despite these discoveries, none of these markers are ubiquitous and specific simultaneously. For example, studies have demonstrated the presence of αSMA^low^ CAFs despite αSMA being previously considered a marker for all activated CAFs (34). More recently, Elyada *et al.* identified PDPN as a robust pan-CAF marker through scRNA-seq, but PDPN is also expressed on lymphatic endothelial cells as well (10, 40). This ultimately results in CAFs being defined in practice by morphology, spatial distribution, and the lack of markers for leukocytes, epithelial, and endothelial cells (32).

In their original study, Ohlünd *et al.* identified (FAP+αSMA+) myCAFs and (αSMA^Low^IL-6^High^) iCAFs as distinct populations through a combination of immunofluorescence and co-culture experiments (34). To isolate for CAFs from mice samples, Ohlünd *et al.* used a combination of outgrowth and clonal isolation. They also performed fluorescence-activated cell sorting (FACS) by staining dissociated single cells with anti-CD45, anti-CD31, and anti-CD140a (PDGFRα), with the PDGFRα+ population representing CAFs and subsequently sorted for RT-qPCR analysis.

On the other hand, Hosein *et al.* profiled the cellular heterogeneity in different stages of PDAC of several genetically engineered mouse models by performing scRNA-seq following dissociation of the whole tumor into single cells without enrichment for CAFs (36). After unsupervised clustering, they identified three clusters of fibroblasts in the normal pancreas and early *KRAS^LSL-G12D/+^Inhk4a^fl/fl^Ptf1a^Cre/+^*(*KIC*) sample. In the late *KIC*, *KPC*, and *KPfC* (*KRAS^LSL-G12D/+^Trp53^fl/fl^Pdx1^Cre/+^*) models, only two clusters were identified, corroborating the myCAF and iCAF populations previously described. However, these two CAF clusters represented only a small fraction of the total cells within their respective samples, suggesting that CAFs may possibly be underrepresented in the scRNA-seq of whole tumor (36). Using a similar method for our orthotopic model, the CAF cluster in the KPC-4545 unenriched sample consisted of other epithelial cells and granulocytes possibly due to overlapping transcriptomic profiles. Furthermore, only one cluster of CAFs was identified in the KPC-3403 unenriched sample, making it challenging to investigate the phenotypic heterogeneity of CAFs within our models. These results highlight the difficulty of studying CAFs specifically in unenriched scRNA-seq, As such, while scRNA-seq has undoubtedly revolutionized the study of CAF subpopulations, challenges remain in capturing more granular details. CAFs are often embedded within the ECM, making them difficult to capture and thus often underrepresented in scRNA-seq of whole tumors, compounding the restraints of limited samples or cell numbers (18). This has led to the development of different protocols to isolate fibroblast-enriched fraction from the whole tumor before scRNA-seq. In their landmark study, of which the methods for this study were based on, Elyada *et al.* enriched for fibroblasts by using antibodies against CD45, CD31, and EPCAM to deplete immune, endothelial, and epithelial cells respectively after dissociating tumors into single cells to obtain the fibroblast enriched fraction (10). This negative selection and sequencing allowed them to corroborate their previous findings of myCAFs and iCAFs while also identifying apCAFs in the mice KPC TME. Similarly, in the data presented in this current study, we were also able to better understand the phenotypic distribution of CAFs in our KPC orthotopic models by conducting scRNA-seq on fibroblast enriched fractions.

Other studies have built upon these phenotypic definitions and sorting methods to isolate specific subpopulations for studying. In their study of apCAFs, Huang *et al.* isolated different subpopulations of CAFs based on the following criteria through FACS: apCAFS (PDPN+MHCII+), iCAFs (PDPN+MHCII-Ly6C+), and myCAFs (PDPN+MHCII-Ly6C+) (37).

In another study, Mucciolo *et al.* sought to further investigate heterogeneity within the myCAF phenotype that express differing levels of CD90 (41). They isolated myCAFs (DAPI-CD31-CD45-EPCAM-PDPN+Ly6C-MHCII-) that were either CD90+ and CD90- and found that the two populations responded differently to EGFR/ERBB2 inhibition through bulk RNA-sequencing. While the discovery of subtype-specific markers like these will allow researchers to better understand the function of different CAF phenotypes, several challenges remain for these sorting experiments. Different dissociation and isolation protocols could lead to artifacts in which transcriptomic profiles are associated with the dissociation method used (42). Furthermore, enriching for CAFs prior to scRNA-seq results in the loss of other cells within the TME, which could have been used to infer cell-cell interactions that are targetable or altered after treatment.

The challenges of identifying CAF subpopulations at a more granular level while not sacrificing the valuable information from other cells within the TME could be addressed through techniques such as single-nuclear RNA sequencing (snRNA-seq) (43). snRNA-seq, which is also compatible with frozen samples, has been shown to have a higher recovery rate of malignant and stromal cells (44, 45). Hwang *et al.* analyzed a snRNA-seq data set consisting of primary PDAC specimen from 43 patients with (n=25) or without (n=18) neoadjuvant treatment prior to surgical resection (46). Through consensus non-negative matrix factorization (cNMF), they identified four recurrent CAF expression programs across the samples, including the myofibroblastic progenitor, immunomodulatory, neurotropic, and adhesive programs. The myofibroblastic program resembles the published myCAF signature while the remaining three overlapped with the published iCAF signature. This demonstrates snRNA-seq’s ability to capture and distinguish different CAF subpopulations in addition to other stromal and tumor cells.

Cellular indexing of transcriptomics and epitopes by sequencing (CITE-seq) can also be further used to unravel the functional heterogeneity of CAFs (47). CITE-seq combines barcoded antibody-based detection of protein markers with transcriptome profiling at the single-cell level (48, 49). Transcriptomic profiling by scRNA-seq alone, while powerful, offers an incomplete picture without information on protein expression and post-translational modifications. And while surface markers can be assessed with multi-color FACS, its targets are limited compared to CITE-seq (47). Given the importance of protein markers in addition to gene expression profile in CAF functional heterogeneity, CITE-seq can be effectively used to study CAFs along with other cells within the TME.

## CONCLUSION

Ultimately, our results demonstrate that isolating a CAF-enriched fraction in our KPC orthotopic tumors captures finer details of CAF heterogeneity through scRNA-seq compared to unenriched single cell dissociates of the whole tumor, but at the cost of other cells within the TME. This warrants the use of other methods such as snRNA-seq and CITE-seq to study the complex cell-cell interaction between CAFs and other cells within the TME and identify how distinct CAF populations respond differently to treatment.

## SUPPLEMENTAL INFORMATION

**Supplementary Table 1.**
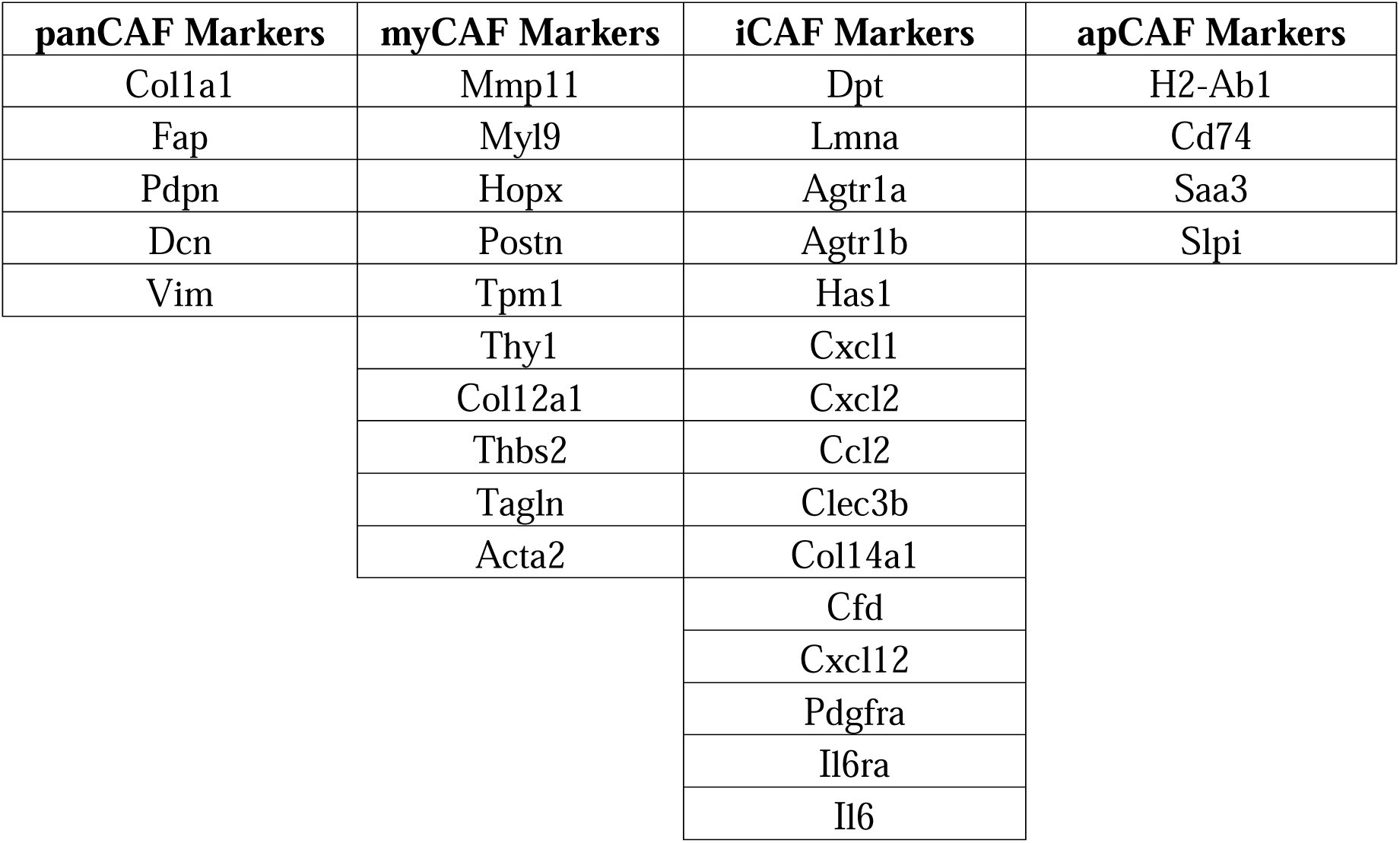
Markers for CAF Phenotypes.

**Supplementary Table 2.**
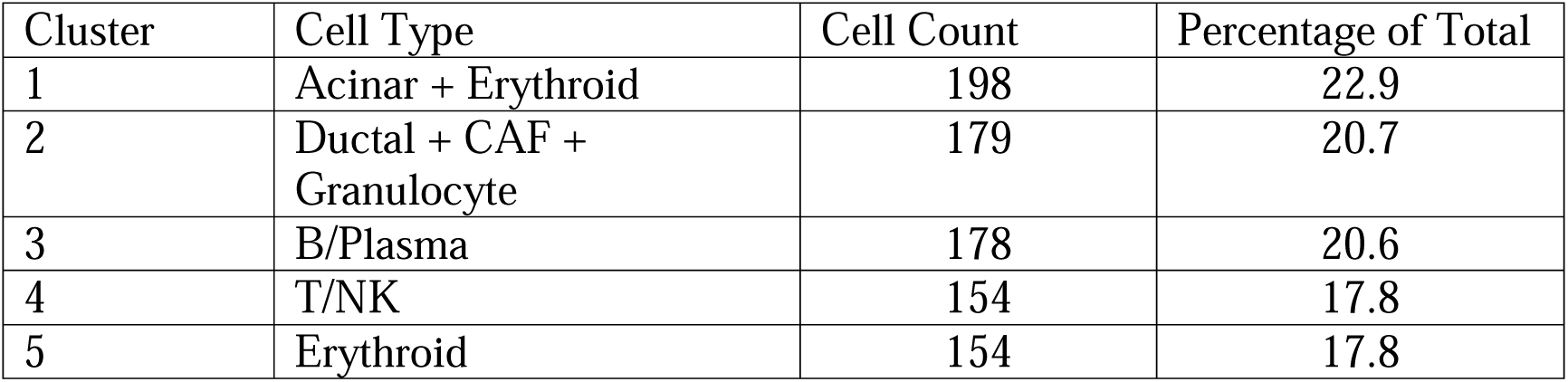
Cell Type Distribution in KPC-4545 Unenriched Sample.

**Supplementary Table 3.**
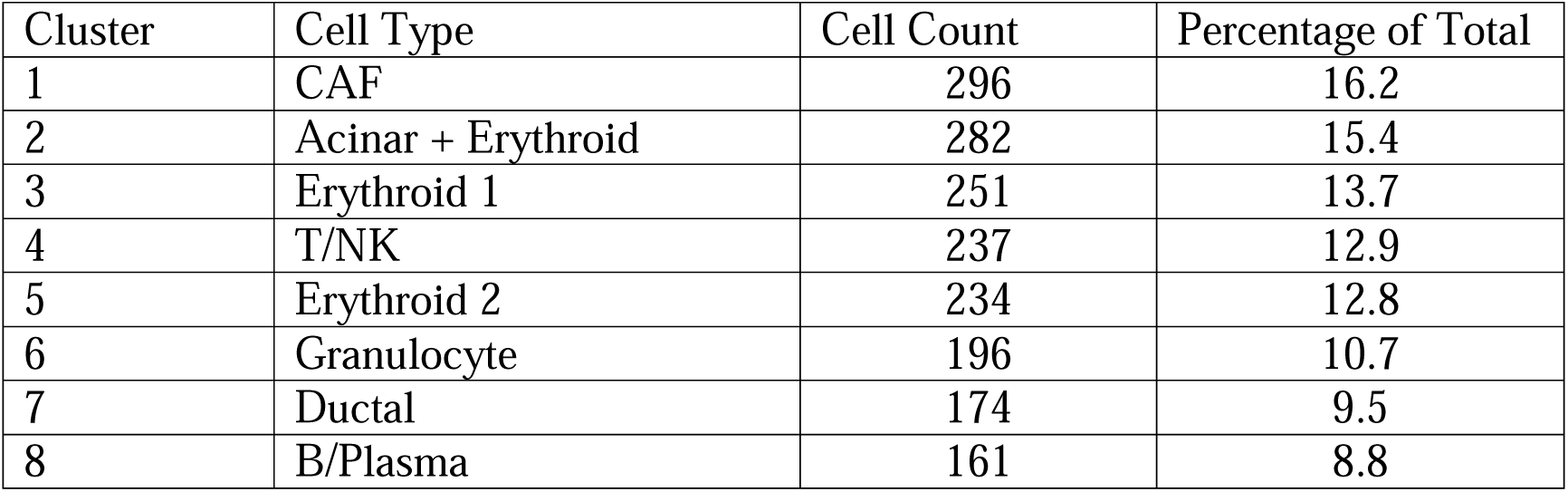
Cell Type Distribution in KPC-3403 Unenriched Sample.

