## Supplementary material for "Comparison of Different Approaches to Single Cell RNA Sequencing of Cancer Associated Fibroblasts": Table S1

**SUPPLEMENTAL INFORMATION**

**Supplementary Table 1.** Markers for CAF Phenotypes

| **panCAF Markers** | **myCAF Markers** | **iCAF Markers** | **apCAF Markers** |
| --- | --- | --- | --- |
| Col1a1 | Mmp11 | Dpt | H2-Ab1 |
| Fap | Myl9 | Lmna | Cd74 |
| Pdpn | Hopx | Agtr1a | Saa3 |
| Dcn | Postn | Agtr1b | Slpi |
| Vim | Tpm1 | Has1 |  |
|  | Thy1 | Cxcl1 |  |
|  | Col12a1 | Cxcl2 |  |
|  | Thbs2 | Ccl2 |  |
|  | Tagln | Clec3b |  |
|  | Acta2 | Col14a1 |  |
|  |  | Cfd |  |
|  |  | Cxcl12 |  |
|  |  | Pdgfra |  |
|  |  | Il6ra |  |
|  |  | Il6 |  |
