## Supplementary material for "Comparison of Different Approaches to Single Cell RNA Sequencing of Cancer Associated Fibroblasts": Table S2

**SUPPLEMENTAL INFORMATION**

**Supplementary Table 2.** Cell Type Distribution in KPC-4545 Unenriched Sample

| Cluster | Cell Type | Cell Count | Percentage of Total |
| --- | --- | --- | --- |
| 1 | Acinar + Erythroid | 198 | 22.9 |
| 2 | Ductal + CAF + Granulocyte | 179 | 20.7 |
| 3 | B/Plasma | 178 | 20.6 |
| 4 | T/NK | 154 | 17.8 |
| 5 | Erythroid | 154 | 17.8 |
