## Supplementary material for "Comparison of Different Approaches to Single Cell RNA Sequencing of Cancer Associated Fibroblasts": Table S3

**SUPPLEMENTAL INFORMATION**

**Supplementary Table 3.** Cell Type Distribution in KPC-3403 Unenriched Sample

| Cluster | Cell Type | Cell Count | Percentage of Total |
| --- | --- | --- | --- |
| 1 | CAF | 296 | 16.2 |
| 2 | Acinar + Erythroid | 282 | 15.4 |
| 3 | Erythroid 1 | 251 | 13.7 |
| 4 | T/NK | 237 | 12.9 |
| 5 | Erythroid 2 | 234 | 12.8 |
| 6 | Granulocyte | 196 | 10.7 |
| 7 | Ductal | 174 | 9.5 |
| 8 | B/Plasma | 161 | 8.8 |
